# The impact of ZIKV infection on gene expression in neural cells over time

**DOI:** 10.1101/2023.08.06.552183

**Authors:** Moreno Magalhães de Souza Rodrigues, Antonio Marques Pereira Júnior, Eduardo Rocha Fukutani, Mariana Araújo-Pereira, Vanessa Riesz Salgado, Artur Trancoso Lopo de Queiroz

**Affiliations:** Fundação Oswaldo Cruz - RO; Center of Data and Knowledge Integration for Health (CIDACS), Instituto Gonçalo Moniz – BA; Multinational Organization Network Sponsoring Translational and Epidemiological Research (MONSTER) Initiative, Salvador, Brazil; União metropolitana de educação e cultura (UNIME), Lauro de Freitas, Bahia, Brazil; Center of Data and Knowledge Integration for Health (CIDACS), Instituto Gonçalo Moniz – BA, Multinational Organization Network Sponsoring Translational and Epidemiological, Research (MONSTER) Initiative, Salvador, Brazil

**Keywords:** ZIKV infection, Neural Stem Cells, Gene expression, Microarray

## Abstract

Zika virus (ZIKV) outbreak caused one of the most significant medical emergencies in the Americas due to associated microcephaly in newborns. To evaluate the impact of ZIKV infection on neuronal cells over time, we retrieved gene expression data from several ZIKV-infected samples obtained at different time point post-infection (pi). Differential gene expression analysis was applied at each time point, with more differentially expressed genes (DEG) identified at 72h pi. There were 5 DEGs (PLA2G2F, TMEM71, PKD1L2, UBD, and TNFAIP3 genes) across all timepoints, which clearly distinguished between infected and healthy samples. The highest expression levels of all five genes were identified at 72h pi. Taken together, our results indicate that ZIKV infection greatly impacts human neural cells at early times of infection, with peak perturbation observed at 72h pi. Our analysis revealed that all five DEGs, in samples of ZIKV-infected human neural stem cells, remained highly upregulated across the timepoints evaluated. Moreover, despite the pronounced inflammatory host response observed throughout infection, the impact of ZIKV is variable over time. Finally, the five DEGs identified herein play prominent roles in infection, and could serve to guide future investigations into virus-host interaction, as well as constitute targets for therapeutic drug development.

## Introduction

The Zika Virus (ZIKV) outbreak in the Americas captured the world’s attention in 2016, leading to one of the most important medical emergencies in the last decade [1]. ZIKV is a mosquito-borne Flavivirus belonging to the Flaviviridae family and is closely related to other viruses (e.g., West Nile Virus, Dengue virus, and Yellow fever virus) that cause arthropod-borne diseases in humans [2]. Generally, these diseases are transmitted to humans by females of infected mosquitoes belonging to the Aedes genus during blood-feeding [3].

In Brazil, the first confirmed case of ZIKV was reported in 2015 [4], followed by the swift spread of the virus [5]. In 2016, more than 26 countries across the Americas and Europe reported cases of the disease [6]. Most often (in over 80% of ZIKV infections in healthy individuals), disease presentation is an asymptomatic infection or characterized by mild symptoms [7–9]. However, less than one year after the first reported case in Salvador, the capital of the state of Bahia-Brazil, increasing reports of Guillain-Barré syndrome (GBS) was noted [10]. Furthermore, in the state of Pernambuco, prenatal ZIKV exposure was associated with an almost 20-fold increase in the incidence of microcephaly [11]. Currently, ZIKV infection during pregnancy could lead to what is known as Congenital Zika Syndrome (CZS). CZS has been linked to brain abnormalities and/or microcephaly or neural tube defects, eye abnormalities, or consequent central nervous system dysfunction among fetuses or infants in the absence of other evidence of brain abnormalities or microcephaly [12].

Since the 2016 outbreak, the effects of ZIKV infection on gene expression in the host’s brain cells have been extensively studied to elucidate the molecular mechanisms related to CZS neuropathogenesis. A previous study explored the effects of ZIKV infection on mRNA and microRNA (miRNA) expression in astrocytes, aiming to identify important miRNAs and pathways related to viral pathology [13]. The effects of ZIKV infection on human neural stem cells (hNSCs) have also recently been studied [14]. Further investigations established the importance of viral infection in hNSCs with respect to the neuropathogenesis of CZS [15–17], thereby corroborating the previous study’s findings, by identifying several miRNAs related to neurodevelopment and oxidative stress. Other studies have also provided insights about ZIKV’s effects on miRNA expression in hNSCs [18], highlighting miRNAs that mediate the suppression of gene networks related to cell cycle progression and stem cell maintenance. Nonetheless, the pathogenesis of ZIKV in neuronal cells has not been completely elucidated, and an mRNA-focused biosignature is still absent from the current literature.

In order to provide further insights into the effects of ZIKV infection on neuronal stem cells and potentially enhance the body of knowledge on CZS development, an integrated gene expression analysis was performed in publicly available gene expression datasets. A similar approach has been widely employed to identify important genes and molecular pathways with the aim of discovering potential biomarkers in several pathologies, such as endometrial cancer [19], Sickle cell disease [20], and HTLV-1 [21]. Related studies have analyzed Aedes aegypti expression datasets to identify important genes and pathways related to mosquito infection by Dengue, Yellow fever, West Nile, and Zika viruses [22,23]. The present investigation analyzed neuronal stem cell gene expression data among 39 samples of which 30 were ZIKV-infected samples collected at different times post-infection and 9 non infected samples.

## Materials and Methods

### Data acquisition

The flow chart for the study selection for this integrated analysis is shown in Fig 1. We have performed a dataset search on the Gene Expression Omnibus (GEO) repository to assess data about the gene expression of neuronal stem cells infected by ZIKV [24]. 86 GEO datasets were founded; however, 77 records were excluded because the samples were not induced pluripotent stem cell-derived human neural stem cells, resulting in 9 datasets. Of these, seven were excluded because they do not have more than one timepoint or because they only have information on non-coding RNAs. Thus, only two datasets were included (GSE97872 and GSE157530).

**Fig 1.**
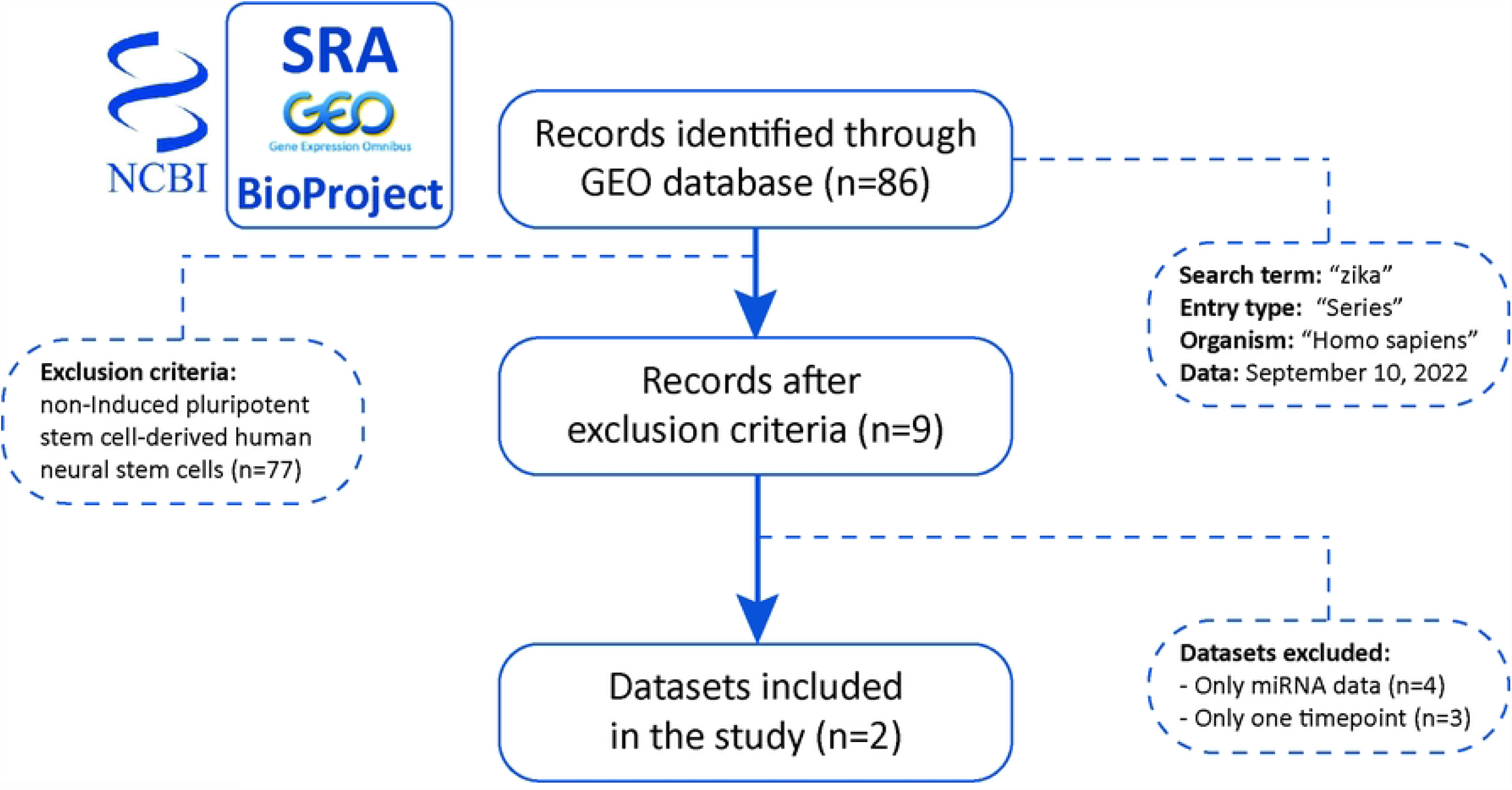
Gene expression data sampling flow chart. Sampling process showing the query terms in the database, the exclusion criteria and the final number of datasets used in the downstream analysis.

The data from both datasets were downloaded from the GEO repository [24] using the GEOquery package [25]. Given that these are public datasets, Ethics Committees authorizations are not required for this paper. A total of 39 samples from human neural stem cells were retrieved, comprising 9 uninfected samples (healthy controls), 6 ZIKV-infected samples collected at 24 hours post-infection (pi), 6 samples at 48h pi, 6 samples at 72h pi, 3 samples at 5 weeks pi, 3 samples at 8 weeks pi,3 samples at 11 weeks pi, and 3 samples at 14 weeks pi. The raw data were log2-transformed and batch-effect corrected using the sva package [26] and then subjected to downstream analysis.

### Differentially expressed genes

Differential expression analysis was employed to compare gene expression profiles in ZIKV-infected hNSCs over time with healthy controls in order to identify differentially expressed genes (DEGs). This analysis was performed using the limma package [27] in R 4.1.0. DEGs were identified using a log fold-change threshold of ±1, with a false discovery rate corrected p-value < 0.05. Comparisons of DEGs at different sample collection times were performed by constructing Venn diagrams using the VennDiagram package [28]. The compareCluster package [29] was then used to scan the REACTOME pathway database [30] for the obtained DEGs to identify enriched pathways at each selected timepoint.

### Evaluation of sample heterogeneity over time

Variation among samples within and between timepoints was evaluated using the molecular degree of perturbation (MDP) package [31], applied to gene expression values following batch correction. All comparisons between timepoints were performed with the Kruskal-Wallis test, followed by Tukey’s posttest.

### Data visualization and dimensionality reduction

To evaluate sample clustering and classification over time, we performed one-sided, unsupervised Ward’s method hierarchical cluster analysis [32], heatmaps [33] and principal component analysis (PCA) using the obtained gene expression values. This approach allowed for the visualization of sample dispersion across timepoints.

## Results

Comparisons between ZIKV-infected and healthy samples revealed a variety of DEGs at different timepoints: 154 DEGS at 24h, 76 at 48h, 299 at 72h, 123 at 5 weeks, 40 at 8 weeks, 19 at 11 weeks, and 34 at 14 weeks (Supplementary Table 1). The most DEGs were identified at the 72h pi timepoint (n=209), in contrast to the fewest (n=1) at 11 weeks pi (Fig 2). A total of five DEGs were shared at all timepoints, genes PLA2G2F, TMEM71, PKD1L2, UBD and TNFAIP3.

**Fig 2.**
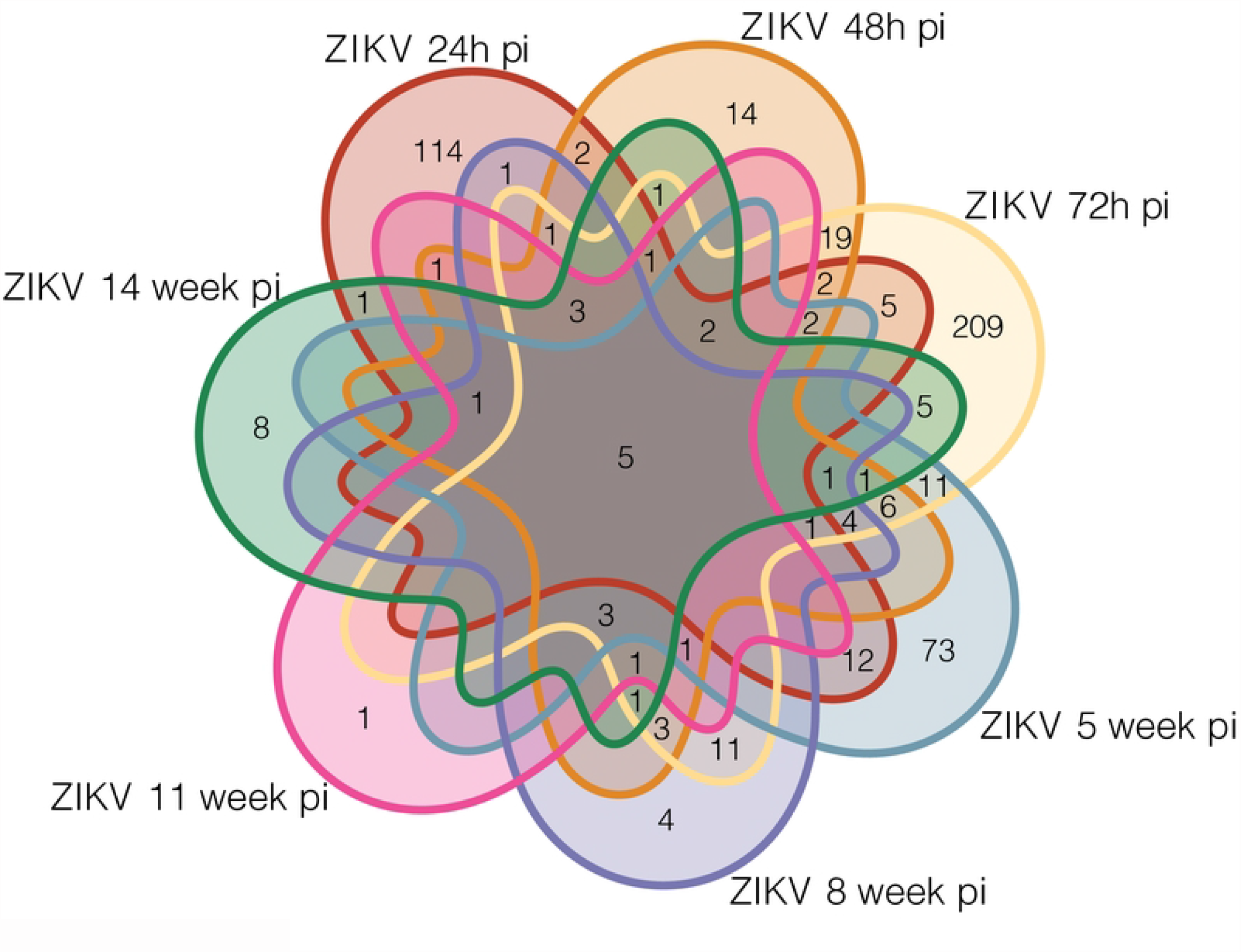
Profile of ZIKV-infected and healthy samples. Venn diagram depicting differentially expressed genes (DEGs) in ZIKV-infected samples collected at different timepoints (comparisons with healthy samples).

We then investigated the pathways associated with specific DEGs identified at each timepoint. This analysis revealed an intense inflammatory response via the enriched pathways Interferon alpha/beta signaling and Interferon Signaling at 48h pi, which persisted until 14 weeks (Fig 2). At 24h pi, the enriched pathways Extracellular matrix organization, Integrin cell surface interactions, Immunoregulatory interactions between a Lymphoid and a non−Lymphoid cell, and Degradation of the extracellular matrix indicated a cell organization and interaction response. At 72h pi, an intensified immune response was evidenced via the enrichment of the pathways Interleukin−4 and Interleukin−13 signaling, NGF−stimulated transcription, Nuclear Events (kinase and transcription factor activation) and Antigen Presentation: Folding, assembly and peptide loading of class I MHC. At 5 weeks pi, a specific pattern of alterations in synapses and cell-signaling was identified via the enriched pathways Regulation of FZD by ubiquitination, Transmission across Chemical Synapses, RUNX2 regulates osteoblast differentiation Trafficking of AMPA receptors, Glutamate binding, activation of AMPA receptors and synaptic plasticity and RUNX2 regulates bone development. However, at 8 weeks pi, an unspecific pattern related to the enrichment of pathways ATF4 activates genes in response to endoplasmic reticulum stress, PERK regulates gene expression, Unfolded Protein Response (UPR) and Response of EIF2AK4 (GCN2) to amino acid deficiency was identified. At 11 weeks pi, just one pathway was enriched: Ovarian tumor domain proteases (Fig 3).

**Fig 3.**
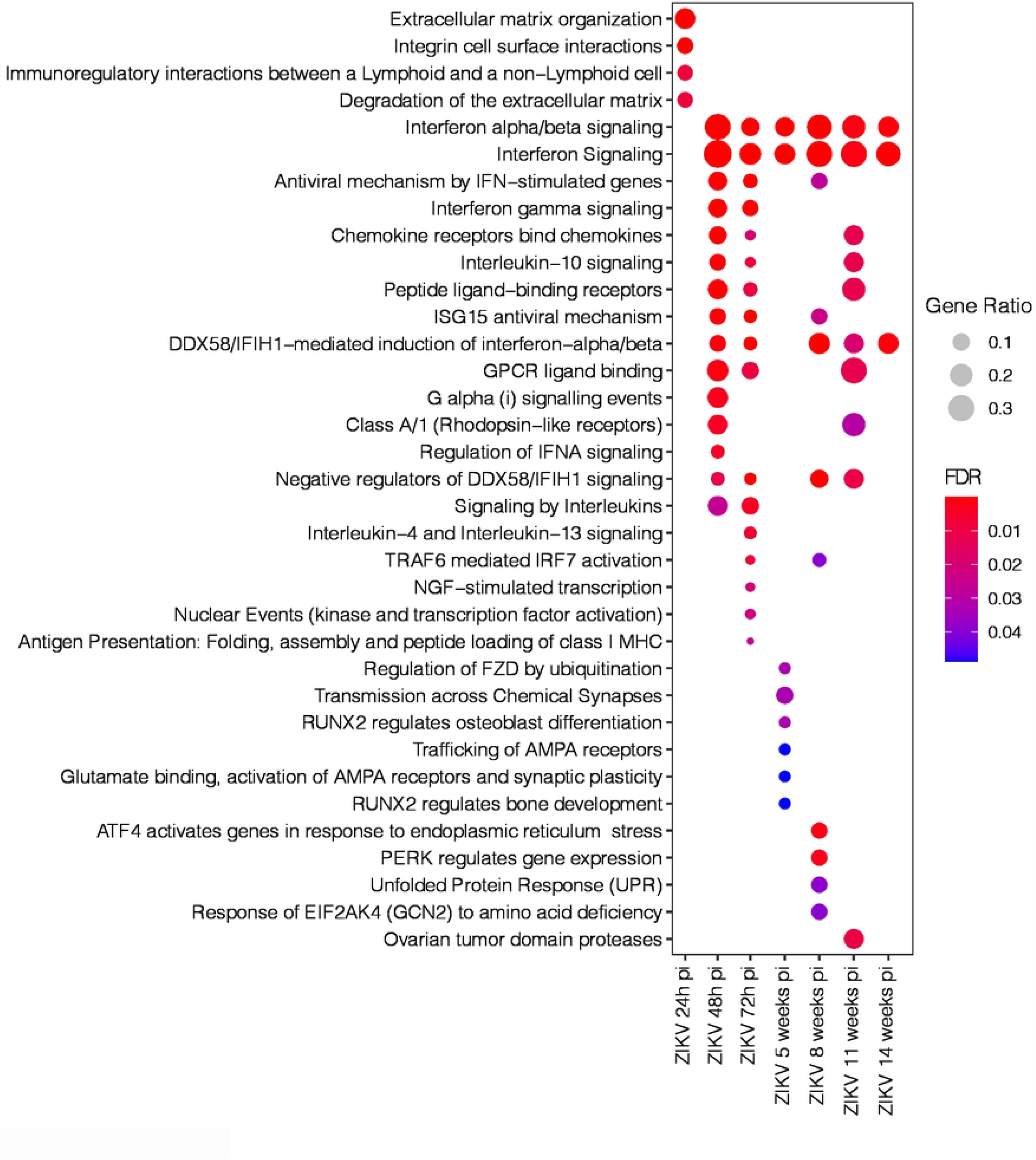
Pathway enrichment analysis involving specific DEGs identified at each sample collection timepoint. Dot size represents pathway gene ratio, while fill colors represent FDR-corrected p-values.

We assessed the sample variations using degree of molecular perturbation and Principal Component Analysis (PCA) in the ZIKV samples to investigate variation across timepoints. We observed variation in ZIKV-infection-associated gene expression at all pi timepoints investigated. The highest variation was identified at 72h pi, significantly greater than at all other timepoints except week 5 (Fig 4A-B), suggesting that ZIKV may exert a stronger impact 72 hours after being bitten by an infected mosquito, with a progressive reduction in intensity over time.

**Fig 4.**
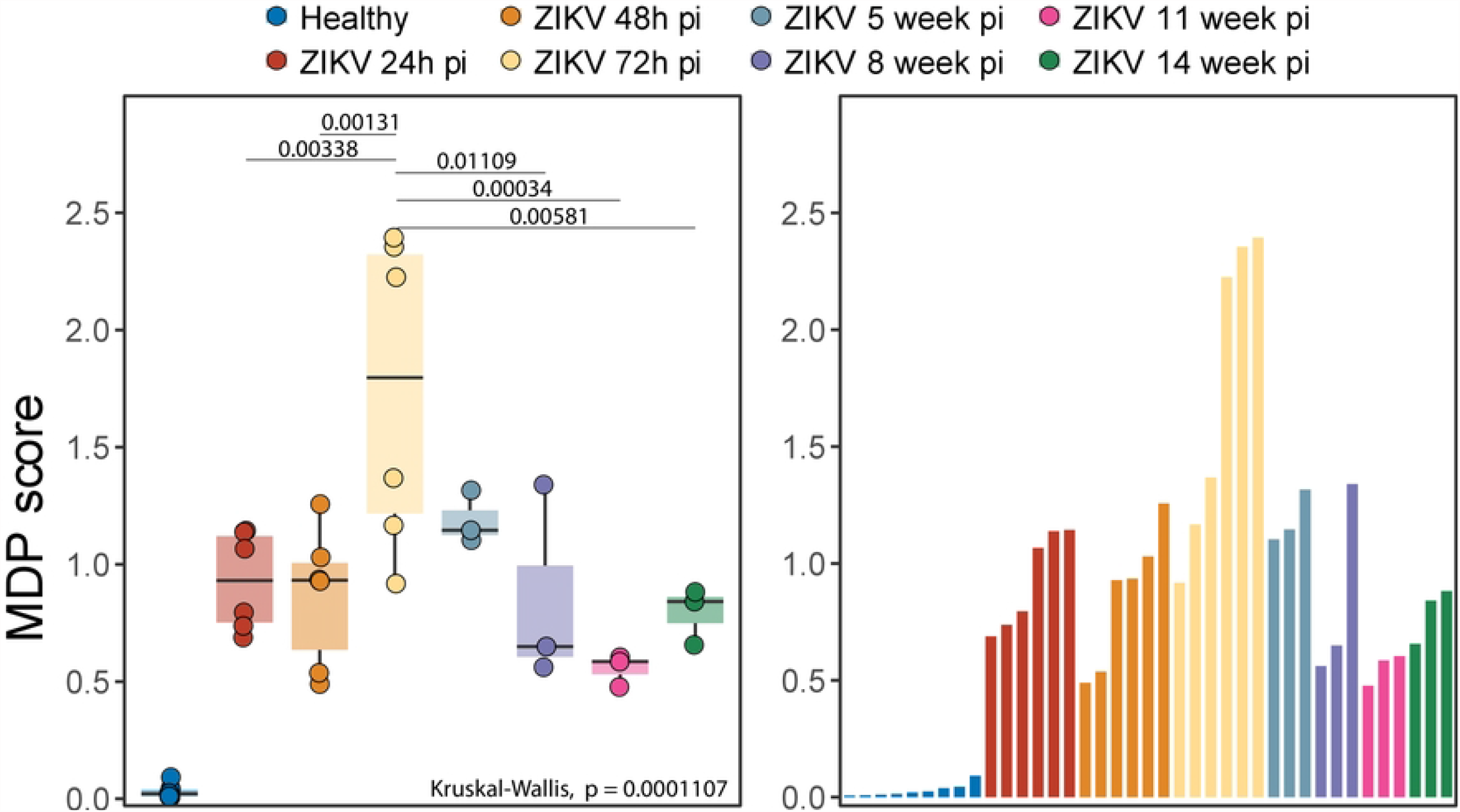
Variation analysis of ZIKV: Molecular Degree of Perturbation analysis in the ZIKV infected samples using healthy samples as baseline. (A) Box Plot of MDP scores over time. Medians were tested using Kruskal-Wallis with Turkey’s post-hoc test. (B) Bar plot represent the value observed in each sample.

The five genes that were significantly differentially expressed at all timepoints were investigated. All were found to be upregulated across the timepoints, with the highest log2 fold-change (log2FC ∼2) seen in PKD1L2, while the lowest was in UBD (log2FC ∼1). Despite the absence of time-specific clustering, heatmap analysis indicated the five genes were capable of differentiating between ZIKV-infected and healthy samples (Figure 5A). Moreover, PCA was applied using these five genes, with specific sample clustering identified at 72h and 48h pi (Fig 5B). We further investigated levels of gene expression of each of the five DEGs across time (Fig 5C). Similar levels of gene expression were found for all five genes, with progressive increases observed at 24h pi and 48h pi, respective peaks at 72h pi, followed by dramatically lower expression levels after 5 weeks pi.

**Fig 5.**
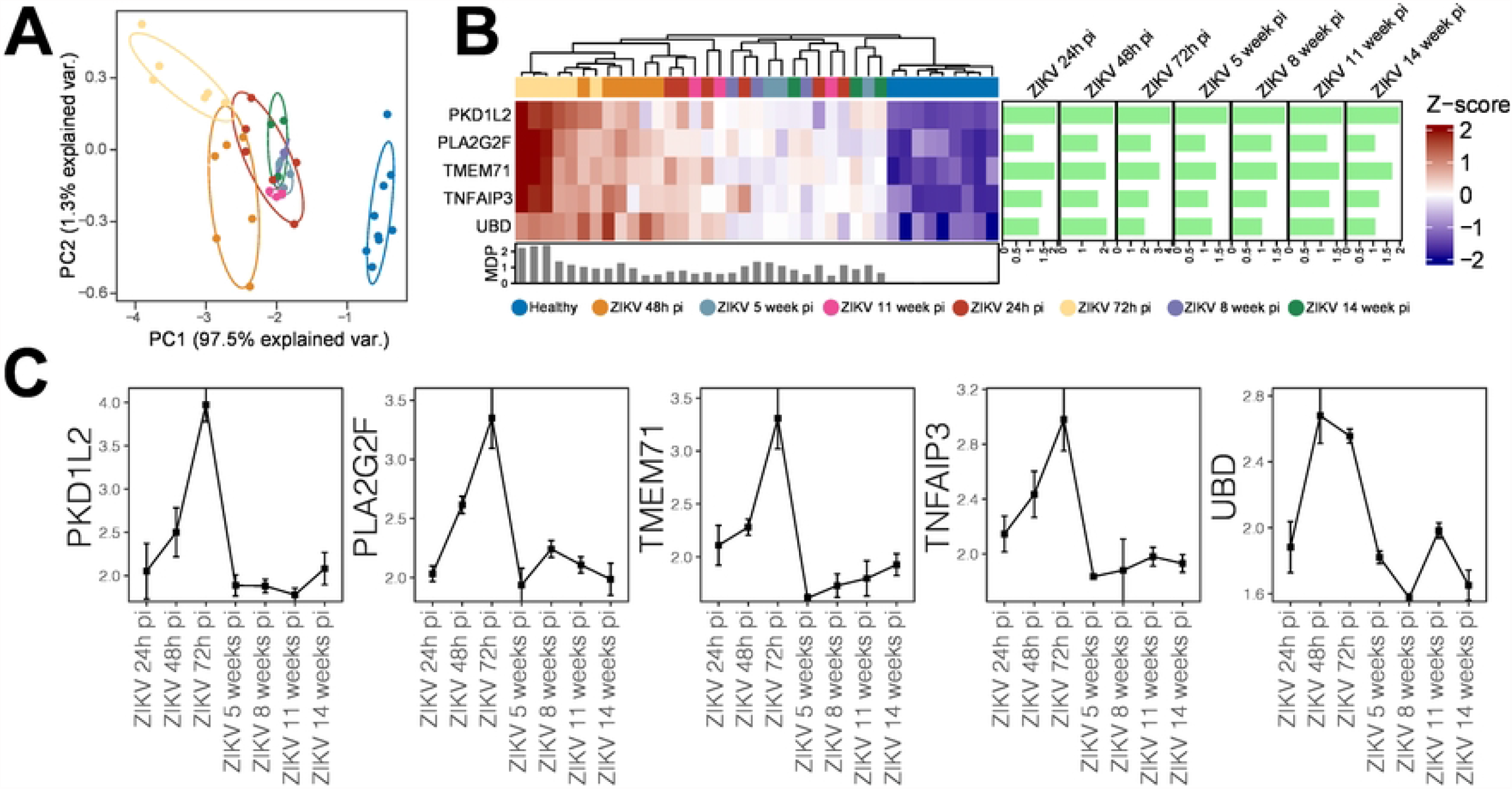
Gene expression levels over time. (A) Heatmap generated from gene expression levels of five DEGs identified at all time points. The colored line (above gene names) reflects the grouping of samples according to gene expression level. The sidebar plot depicts significant log2 fold-change values (FDR<0.05) for each gene across the evaluated timepoints compared to healthy controls. The bottom bar plot reflects the MDP score values of each sample. (B) Gene expression levels of 5 shared DEGs across the evaluated timepoints: Squares indicate median expression levels at each respective timepoint, with bars representing standard error.

## Discussion

Both human and animal studies have detected ZIKV in neuronal cells, including progenitor and mature stem cells, as well as astrocytes in vitro. The virus has also been detected in neurosensory tissue, such as the retina and optic nerve [34]. Research has shown the alterations viruses cause in cellular pathways create optimal environments for their replication. These strategies make energy resources available for viruses, provide substrate for viral particles and increase the survival of infected cells to consequently ensure viral replication [35]. Accordingly, enhancing our understanding of the metabolic alterations induced by viruses may yield useful insights for therapeutic applications in viral diseases. The exact mechanisms (biochemical and cellular) by which the neuropathogenesis of ZIKV happens remains poorly understood [36].

Recent in vitro studies have demonstrated that ZIKV avoids the IFN response (pathway associated with IFN-stimulated response) by destroying STAT2 via proteasomal degradation. The present study conducted transcriptomic analysis on neural cells infected with ZIKV at several post-infection timepoints in order to identify not only differentially expressed genes in comparison to uninfected controls, but also across different time intervals. We further investigated pathways associated with infection at each time interval. In addition to molecular/biochemical alterations caused by ZIKV in host cells on a transcriptomic level, these can also be observed at the proteomic level. Another study investigating the proteome involved in ZIKV infection in human mesenchymal stem cells (hMSC) observed that over 100 proteins were altered by viral infection, many known to be related to different neuropathologies [37]. A systematic review of several studies investigating ZIKV-infected organoids identified that viral load was increased after 72 hours regardless of ZIKV viral lineage, a finding consistent with our data that indicates a peak in DEG detection at 72 hours post-infection [37,38].

The present study identified five DEGs shared across all seven-time intervals evaluated: PLA2G2F, TMEM71, PKD1L2, UBD, and TNFAIP3. Another study investigating the effect of ZIKV infection on NSC (neural stem cells) in miRNA signatures and gene expression reported alterations in several signaling pathways due to ZIKV [14]. Similarly to the present study, these authors observed increased TMEM71 expression (2.5-, 8.7-, and 35.8-fold increases respectively after 24, 48, and 72 hours of infection) and PKD1L2 (2.8-, 9.6-, and 49.9-fold increases at 24, 48, and 72 after infection, respectively). The authors further reported the dysregulation of several pathways involved in nervous system development and cell cycle, and associated these dysregulations with the formation of microcephaly. Finally, a study conducted in non-human primates infected with ZIKV observed an up-regulation of genes related to interferon, which stands in agreement with the present results [39].

In another study that used NPCs (neuronal precursor cells) to investigate the effect of ZIKV infection on cellular pathways, the authors attempted to establish correlations with clinical aspects of disease (microcephaly, epilepsy, etc.), observing that viral infection activates a variety of inflammatory signaling, similar to our observations after 48h in affected pathways [40]. Furthermore, in 2017, a group of researchers investigated the presence of ZIKV in the cerebrospinal fluid (CSF) and lymph nodes (LN) in rhesus monkeys. These authors detected upregulation in the mTOR, proinflammatory, and anti-apoptotic signaling pathways, with the simultaneous downregulation of extracellular matrix signaling pathways. They reported the strong induction of IFN-alpha and other antiviral pathways at 2, 4 and 6 days post-infection, as well as the upregulation of inflammasome components, cytokines and immunomodulatory pathways [41].

In conclusion, the present findings provide evidence that ZIKV infection in human neuronal cells induced the differential expression of several genes that varied across a range of time intervals post-infection. MDP analysis revealed the 72h pi timepoint as significant for heterogenous gene expression, and the DEGs identified were linked to interleukin and extracellular matrix signaling pathways. Moreover, five DEGs observed across all timepoints were found to be capable of distinguishing between ZIKV-infected and healthy hNSCs. Taken together, those genes could serve as a workbench for future works using RNA-seq data. This methodology allows us to estimate both host and viral RNA expression. This information together would provide useful insights into the viral/host relationship and cell damage process of neuropathogenesis.

## Supporting information

**S1 Table. Multiple comparison of genes**.

## Data availability

Data used in this project are free available at Gene Expression Omnibus (GEO) repository.

## Acknowledgments

The authors would like to thank Andris K. Walter for critical analysis, English language revision and manuscript copyediting assistance.

## Author Contributions

**Conceptualization:** Eduardo Fukutani, Vanessa Salgado, Antonio Júnior, Moreno Rodrigues and Artur Queiroz.

**Data curation:** Antonio Júnior; Moreno Rodrigues.

**Formal analysis:** Artur Queiroz.

**Funding acquisition:** Artur Queiroz.

**Investigation:** Artur Queiroz.

**Methodology:** Eduardo Fukutani and Artur Queiroz.

**Project administration:** Moreno Rodrigues.

**Resources:** Moreno Rodrigues.

**Writing – original draft:** Antonio Júnior, Eduardo Fukutani, Vanessa Salgado, Moreno Rodrigues and Artur Queiroz.

**Writing – review & editing:** Antonio Júnior, Eduardo Fukutani, Vanessa Salgado, Moreno Rodrigues and Artur Queiroz.

## References

1. Chang C, Ortiz K, Ansari A, Gershwin ME. The Zika outbreak of the 21st century. J Autoimmun. 2016; 1–13. doi:10.1016/j.jaut.2016.02.006

2. Choumet V, Despres P. La dengue et autres infections à flavivirus. Rev Sci Tech l’OIE. 2015;34: 467–478. doi:10.20506/rst.34.2.2372

3. Gregory CJ, Oduyebo T, Brault AC, Brooks JT, Chung KW, Hills S, et al. Modes of Transmission of Zika Virus. J Infect Dis. 2017;216: S875–S883. doi:10.1093/infdis/jix396

4. Zanluca C, Melo VCA, Mosimann ALP, Santos GI V., Santos CND, Luz K. First report of autochthonous transmission of Zika virus in Brazil. Mem Inst Oswaldo Cruz. 2015;110: 569–572. doi:10.1590/0074-02760150192

5. Organization WH. Zika situation report: neurological syndrome and congenital anomalies. 2016.

6. Zammarchi L, Tappe D, Fortuna C, Remoli ME, Günther S, Venturi G, et al. Zika virus infection in a traveller returning to Europe from Brazil, March 2015. Eurosurveillance. 2015;20: 21153. doi:10.2807/1560-7917.ES2015.20.23.21153

7. Carteaux G, Maquart M, Bedet A, Contou D, Brugières P, Fourati S, et al. Zika Virus Associated with Meningoencephalitis. N Engl J Med. 2016;374: 1595–1596. doi:10.1056/NEJMc1602964

8. Mlakar J, Korva M, Tul N, Popović M, Poljšak-Prijatelj M, Mraz J, et al. Zika Virus Associated with Microcephaly. N Engl J Med. 2016;374: 951–958. doi:10.1056/NEJMoa1600651

9. Cao-Lormeau VM, Blake A, Mons S, Lastère S, Roche C, Vanhomwegen J, et al. Guillain-Barré Syndrome outbreak associated with Zika virus infection in French Polynesia: a case-control study. Lancet. 2016;387: 1531–1539. doi:10.1016/S0140-6736(16)00562-6

10. Styczynski AR, Malta JMAS, Krow-Lucal ER, Percio J, Nóbrega ME, Vargas A, et al. Increased rates of Guillain-Barré syndrome associated with Zika virus outbreak in the Salvador metropolitan area, Brazil. Carvalho MS, editor. PLoS Negl Trop Dis. 2017;11: e0005869. doi:10.1371/journal.pntd.0005869

11. Fauci AS, Morens DM. Zika Virus in the Americas — Yet Another Arbovirus Threat. N Engl J Med. 2016;374: 601–604. doi:10.1056/NEJMp1600297

12. Freitas DA, Souza-Santos R, Carvalho LMA, Barros WB, Neves LM, Brasil P, et al. Congenital Zika syndrome: A systematic review. Fujioka K, editor. PLoS One. 2020;15: e0242367. doi:10.1371/journal.pone.0242367

13. Kozak R, Majer A, Biondi M, Medina S, Goneau L, Sajesh B, et al. MicroRNA and mRNA Dysregulation in Astrocytes Infected with Zika Virus. Viruses. 2017;9: 297. doi:10.3390/v9100297

14. Tabari D, Scholl C, Steffens M, Weickhardt S, Elgner F, Bender D, et al. Impact of Zika Virus Infection on Human Neural Stem Cell MicroRNA Signatures. Viruses. 2020;12: 1219. doi:10.3390/v12111219

15. Dang J, Tiwari SK, Lichinchi G, Qin Y, Patil VS, Eroshkin AM, et al. Zika Virus Depletes Neural Progenitors in Human Cerebral Organoids through Activation of the Innate Immune Receptor TLR3. Cell Stem Cell. 2016;19: 258–265. doi:10.1016/j.stem.2016.04.014

16. Li C, Xu D, Ye Q, Hong S, Jiang Y, Liu X, et al. Zika Virus Disrupts Neural Progenitor Development and Leads to Microcephaly in Mice. Cell Stem Cell. 2016;19: 120–126. doi:10.1016/j.stem.2016.04.017

17. Tang H, Hammack C, Ogden SC, Wen Z, Qian X, Li Y, et al. Zika Virus Infects Human Cortical Neural Progenitors and Attenuates Their Growth. Cell Stem Cell. 2016;18: 587–590. doi:10.1016/j.stem.2016.02.016

18. Dang JW, Tiwari S., Qin Y, Rana TM. Genome-wide Integrative Analysis of Zika-Virus-Infected Neuronal Stem Cells Reveals Roles for MicroRNAs in Cell Cycle and Stemness. Cell Rep. 2019;27: 3618–3628.e5. doi:10.1016/j.celrep.2019.05.059

19. O’Mara TA, Zhao M, Spurdle AB. Meta-analysis of gene expression studies in endometrial cancer identifies gene expression profiles associated with aggressive disease and patient outcome. Sci Rep. 2016;6: 36677. doi:10.1038/srep36677

20. Hounkpe BW, Fiusa MML, Colella MP, Costa LNG, Benatti RO, Olalla Saad ST, et al. Role of innate immunity-triggered pathways in the pathogenesis of Sickle Cell Disease: a meta-analysis of gene expression studies. Sci Rep. 2015;5: 17822. doi:10.1038/srep17822

21. Fukutani ER, Ramos PIP, Kasprzykowski JI, Azevedo LG, Rodrigues MMS, Lima JVOP, et al. Meta-Analysis of HTLV-1-Infected Patients Identifies CD40LG and GBP2 as Markers of ATLL and HAM/TSP Clinical Status: Two Genes Beat as One. Front Genet. 2019;10. doi:10.3389/fgene.2019.01056

22. Fukutani KF, Kasprzykowski JI, Paschoal AR, Gomes MS, Barral A, de Oliveira CI, et al. Meta-Analysis of Aedes aegypti Expression Datasets: Comparing Virus Infection and Blood-Fed Transcriptomes to Identify Markers of Virus Presence. Front Bioeng Biotechnol. 2018;5. doi:10.3389/fbioe.2017.00084

23. Fukutani E, Rodrigues M, Kasprzykowski JI, Araujo CF, Paschoal AR, Ramos PIP, et al. Follow up of a robust meta-signature to identify Zika virus infection in Aedes aegypti: another brick in the wall. Mem Inst Oswaldo Cruz. 2018;113. doi:10.1590/0074-02760180053

24. Edgar R, Domrachev M, ALsh AE. Gene Expression Omnibus: NCBI gene expression and hybridization array data repository. Nucleic Acids Res. 2002;30: 207–210. doi:10.1093/nar/30.1.207

25. Davis S, Meltzer PS. GEOquery: a bridge between the Gene Expression Omnibus (GEO) and BioConductor. Bioinformatics. 2007;23: 1846–1847. doi:10.1093/bioinformatics/btm254

26. Leek JT, Johnson WE, Parker HS, Jaffe AE, Storey JD. The sva package for removing batch effects and other unwanted variation in high-throughput experiments. Bioinformatics. 2012;28: 882–883. doi:10.1093/bioinformatics/bts034

27. Ritchie ME, Phipson B, Wu D, Hu Y, Law CW, Shi W, et al. limma powers differential expression analyses for RNA-sequencing and microarray studies. Nucleic Acids Res. 2015;43: e47–e47. doi:10.1093/nar/gkv007

28. Chen H, Boutros PC. VennDiagram: a package for the generation of highly-customizable Venn and Euler diagrams in R. BMC Bioinformatics. 2011;12: 35. doi:10.1186/1471-2105-12-35

29. Yu G, Wang LG, Han Y, He QY. clusterProfiler: an R Package for Comparing Biological Themes Among Gene Clusters. Omi A J Integr Biol. 2012;16: 284–287. doi:10.1089/omi.2011.0118

30. Yu G, He QY. ReactomePA: an R/Bioconductor package for reactome pathway analysis and visualization. Mol Biosyst. 2016;12: 477–479. doi:10.1039/C5MB00663E

31. Oliveira-de-Souza D, Vinhaes CL, Arriaga MB, Kumar NP, Cubillos-Angulo JM, Shi R, et al. Molecular degree of perturbation of plasma inflammatory markers associated with tuberculosis reveals distinct disease profiles between Indian and Chinese populations. Sci Rep. 2019;9: 8002. doi:10.1038/s41598-019-44513-8

32. Ward JH. Hierarchical Grouping to Optimize an Objective Function. J Am Stat Assoc. 1963;58: 236–244. doi:10.1080/01621459.1963.10500845

33. Gu Z, Eils R, Schlesner M. Complex heatmaps reveal patterns and correlations in multidimensional genomic data. Bioinformatics. 2016;32: 2847–2849. doi:10.1093/bioinformatics/btw313

34. Miner JJ, Diamond MS. Zika Virus Pathogenesis and Tissue Tropism. Cell Host Microbe. 2017;21: 134–142. doi:10.1016/j.chom.2017.01.004

35. Sanchez EL, Lagunoff M. Viral activation of cellular metabolism. Virology. 2015;479–480: 609–618. doi:10.1016/j.virol.2015.02.038

36. Acosta-Ampudia Y, Monsalve DM, Castillo-Medina LF, Rodríguez Y, Pacheco Y, Halstead S, et al. Autoimmune Neurological Conditions Associated With Zika Virus Infection. Front Mol Neurosci. 2018;11. doi:10.3389/fnmol.2018.00116

37. Beys-da-Silva WO, Rosa RL, Santi L, Berger M, Park SK, Campos AR, et al. Zika Virus Infection of Human Mesenchymal Stem Cells Promotes Differential Expression of Proteins Linked to Several Neurological Diseases. Mol Neurobiol. 2019;56: 4708–4717. doi:10.1007/s12035-018-1417-x

38. Sutarjono B. Can We Better Understand How Zika Leads to Microcephaly? A Systematic Review of the Effects of the Zika Virus on Human Brain Organoids. J Infect Dis. 2019;219: 734–745. doi:10.1093/infdis/jiy572

39. Chiu CY, Sánchez-San Martín C, Bouquet J, Li T, Yagi S, Tamhankar M, et al. Experimental Zika Virus Inoculation in a New World Monkey Model Reproduces Key Features of the Human Infection. Sci Rep. 2017;7: 17126. doi:10.1038/s41598-017-17067-w

40. Rolfe AJ, Bosco DB, Wang J, Nowakowski RS, Fan J, Ren Y. Bioinformatic analysis reveals the expression of unique transcriptomic signatures in Zika virus infected human neural stem cells. Cell Biosci. 2016;6: 42. doi:10.1186/s13578-016-0110-x

41. Aid M, Abbink P, Larocca RA, Boyd M, Nityanandam R, Nanayakkara O, et al. Zika Virus Persistence in the Central Nervous System and Lymph Nodes of Rhesus Monkeys. Cell. 2017;169: 610–620.e14. doi:10.1016/j.cell.2017.04.008

